# Cell-Type Specific Connectivity of Whisker-Related Sensory and Motor Cortical Input to Dorsal Striatum

**DOI:** 10.1101/2023.03.06.531405

**Authors:** Branden D. Sanabria, Sindhuja S. Baskar, Alex J. Yonk, Christian R. Lee, David J. Margolis

## Abstract

The anterior dorsolateral striatum (DLS) is heavily innervated by convergent excitatory projections from the primary motor (M1) and sensory cortex (S1) and is considered an important site of sensorimotor integration. M1 and S1 corticostriatal synapses have functional differences in the strength of their connections with striatal spiny projection neurons (SPNs) and fast-spiking interneurons (FSIs) in the DLS, and as a result exert an opposing influence on sensory-guided behaviors. In the present study, we tested whether M1 and S1 inputs exhibit differences in the subcellular anatomical distribution onto striatal neurons. We injected adeno-associated viral vectors encoding spaghetti monster fluorescent proteins (sm.FPs) into M1 and S1, and used confocal microscopy to generate 3D reconstructions of corticostriatal inputs to single identified SPNs and FSIs obtained through ex-vivo patch-clamp electrophysiology. We found that SPNs are less innervated by S1 compared to M1, but FSIs receive a similar number of inputs from both M1 and S1. In addition, M1 and S1 inputs were distributed similarly across the proximal, medial, and distal regions of SPNs and FSIs. Notably, clusters of inputs were prevalent in SPNs but not FSIs. Our results suggest that SPNs have stronger functional connectivity to M1 compared to S1 due to a higher density of synaptic inputs. The clustering of M1 and S1 inputs onto SPNs but not FSIs suggest that cortical inputs are integrated through cell-type specific mechanisms and more generally have implications for how sensorimotor integration is performed in the striatum.

**Significance Statement:** The dorsolateral striatum (DLS) is a key brain area involved in sensorimotor integration due to its dense innervation by the primary motor (M1) and sensory cortex (S1). However, the quantity and anatomical distribution of these inputs to the striatal cell population has not been well characterized. In this study we demonstrate that corticostriatal projections from M1 and S1 differentially innervate spiny projection neurons (SPNs) and fast-spiking interneurons (FSIs) in the DLS. S1 inputs innervate SPNs less than M1 and are likely to form synaptic clusters in SPNs but not in FSIs. These findings suggest that sensorimotor integration is partly achieved by differences in the synaptic organization of corticostriatal inputs to local striatal microcircuits.

## Introduction

Sensorimotor integration is the ability to identify important stimuli from the environment and use this information to guide behaviors essential for survival (Machado et al, 2010). Neurons in the primary motor (M1) and primary sensory (S1) cortices play a role in this process by sending excitatory glutamatergic projections to the striatum, a subcortical region, and the main input nucleus of the basal ganglia network (Hoffer & Alloway, 2001). Generally, cortical projections from functionally related brain regions show higher density and significant overlap amongst their projection fields in the striatum (Bolam et al, 2002; Hintiryan et al, 2016; Hunnicutt et al, 2016). Notably, the dorsolateral striatum (DLS) is a site of overlap between inputs from M1 and S1 and plays a critical role in sensorimotor integration (Hoffer & Alloway, 2001; Hunnicutt et al, 2016; Makino et al, 2016; Hooks et al, 2018; Smith et al, 2022), but how these inputs are integrated by different cell types in DLS is still unclear.

The DLS mostly contains inhibitory GABAergic spiny projection neurons (SPNs) and a sparse population of various interneurons (Kawaguchi, 1993 & 1997; Tepper et al, 2018). Activation of SPNs that express the D1 dopamine receptor constitute the “direct” pathway and promote selected motor actions, while SPNs that express the D2 dopamine receptor constitute the “indirect” pathway and inhibit non selected motor actions (Kreitzer & Malenka, 2008). One major interneuron type, the parvalbumin-positive fast spiking interneuron (FSI), is a potent inhibitor of striatal SPNs and involved in sensorimotor transformation of behavior (Lee et al, 2019; Gritton et al, 2019; Martiros et al, 2019). For example, stimulation of S1 inputs to FSIs in the DLS suppresses responding to both rewarded and unrewarded cues in a whisker-based texture discrimination task. In contrast, stimulation of the M1 inputs, which has a different balance of functional innervation to SPNs compared to S1, promotes behavioral responses in the same task (Lee et al, 2019). SPNs and FSIs rely on excitatory inputs along the extent of their elaborate dendrites to drive action potential discharge (Kawaguchi, 1993 & 1997; Stern et al, 1998; Tepper et al, 2018). Therefore, both the local connectivity and external synaptic inputs are important factors of integration in the DLS (Straub et al, 2016; Hjorth et al, 2020).

While M1 and S1 inputs are capable of evoking excitatory postsynaptic potentials (EPSPs) in SPNs and FSIs, recent reports have found that S1 inputs generate larger responses in FSIs compared to SPNs (Lee et al, 2019; Johansson & Silberberg, 2020). Conversely, Lee et al. (2019) found that the response to stimulation of M1 inputs was similar for SPNs and FSIs. Potential differences in synaptic efficacy of M1 and S1 corticostriatal inputs to striatal neurons could reflect presynaptic, postsynaptic, or structural differences (Fino et al, 2005; Perrin & Venance, 2019; Badreddine et al, 2022), mechanisms that could interact to produce differences in functional innervation. Two properties that have been associated with the efficacy of total synaptic input that a cell receives are the number of synaptic inputs and the distribution of synapses along the dendritic arbor (Rall, 1967; Magee, 2000). Understanding how M1 and S1 inputs are organized onto SPNs and FSIs can reveal how the DLS integrates functionally related signals and commands downstream basal ganglia nuclei, which ultimately affect behavior. Here, we investigate the structural input to SPNs and FSIs and provide evidence for the differential innervation of striatal neurons by M1 and S1.

## Materials and Methods

### Animals

All procedures involving animals were approved by the Rutgers University Institutional Animal Care and Use Committee (Protocol #: 999900197). Experiments were performed on wild-type male and female mice on a C57BL/6J background of at least 3 months of age at the time of the first surgical procedures. Mice were housed in a reverse light cycle room (lights off from 08:00 to 20:00) with food and water available *ad libitum*. All mice underwent two simultaneous unilateral Adeno Associated Virus (AAV) injections between 35 and 104 days old and all mice were euthanized after at least three weeks post injection.

### Viral injection

Briefly, mice were anesthetized with isoflurane (4% induction, 0.8%–1.5% maintenance) and placed onto a stereotaxic frame (Kopf) with a feedback-controlled heating blanket maintained at ∼36°C (FHC). Rimadyl (5 mg/kg; Hospira) and Bupivacaine (0.25%, 0.1 mL; Fresenius Kabi) were injected subcutaneously into the left flank and scalp, respectively. The scalp was cleaned with Betadine (Purdue Products) followed by 70% ethanol three times. A midline incision was made, and the skull was exposed and leveled relative to bregma and lambda in the dorsoventral plane. A craniotomy was made above both injection sites, and ∼270 nL of pAAV.CAG.Flex.Ruby2sm-Flag.WPRE.SV40 (#98928; Addgene), pENN.AAV.CAG.Flex.GFPsm_myc.WPRE.SV40 (#98927; Addgene) mixed 1:1 with pAAV.CAMKII.Cre.SV40 (#105558; Addgene), or pAAV.CAG.GFPsm-myc.WPRE.SV40 (#98926; Addgene) diluted 1:1 in 0.1M PBS were pressure injected into either left whisker M1 (anteroposterior (AP): +1.6mm, mediolateral (ML): +1.5mm, dorsoventral (DV): -0.6mm) or left whisker S1 (AP: -1.0mm, ML: +3.3mm, DV: -0.6mm) via a glass micropipette (BlueBrand IntraMark) over the course of 5 minutes followed by another 5 minute delay to permit viral diffusion. After the micropipette was slowly raised, the scalp was closed and secured with sutures and tissue glue. After surgery, mice were placed in clean, temporary housing and monitored for 72 hours. After this monitoring period, mice were transferred to their home cages and allowed to recover for at least 3 weeks to allow for viral expression to fully label neuronal processes.

### Whole cell patch clamp recordings

Mice were induced with 3% isoflurane, deeply anesthetized with ketamine-xylazine (300/30 mg/kg), and transcardially perfused with recovery artificial cerebrospinal fluid (ACSF) containing 103mM NMDG, 2.5mM KCl, 1.2mM NaH_2_PO_4_, 30mM NaHCO_3_, 20mM HEPES, 25mM Glucose, 101mM HCl (1N), 10mM MgSO_4_, 2mM Thiourea, 3mM Sodium Pyruvate, 12mM N-Acetyl-L-Cysteine, and 0.5mM CaCl_2_ (saturated with 95% O_2_ and 5% CO_2_). After decapitation and extraction, the brain was glued to a vibratome stage, immersed in RT oxygenated recovery ACSF, and 300μm coronal sections were cut using a Leica VT1200S vibratome. Sections were immediately transferred to the same oxygenated medium at 35°C for ∼5 min, after which they were transferred to oxygenated external ACSF containing 124mM NaCl, 2.5mM KCl, 26mM NaHCO_3_, 1.2mM NaH_2_PO_4_, 10mM Glucose, 3mM Sodium Pyruvate, 1mM MgCl_2_, and 2mM CaCl_2_ at RT for at least 1 hour before use.

Whole-cell patch clamp recordings were obtained in a chamber that was constantly perfused (2-4 ml/min) with oxygenated external ACSF at 34°C. Sections and neurons were visualized using infrared differential interference contrast (IR-DIC) microscopy with a IR1000 CCD camera (Dage-MTI) mounted onto an BX51-WI upright microscope (Olympus) fitted with two switchable lenses: a 4X air lens and a 40X water-immersion lens. Patch pipettes (2-5MΩ) were fabricated by pulling borosilicate glass micropipettes (2mm o.d., Warner Instruments) via a P-1000 horizontal puller (Sutter Instruments). The internal pipette solution contained 130mM K methanesulfonate, 10mM KCl, 10mM HEPES, 2mM MgCl_2_, 4mM Na_2_ATP, 0.4mM Na_2_GTP at pH 7.25 and 290-295mOsm/L. In addition, 2% biocytin was freshly dissolved in the internal solution on each recording day. Current clamp recordings were obtained from neurons within the anterior dorsal striatum (approximately 1.4 to 0.4mm relative to bregma). Once a selected cell was patched, it was subjected to hyperpolarizing and depolarizing current steps (-500pA to 500pA, 100pA steps, 500ms, 11 sweeps) for post-hoc electrophysiological analysis and identification. Additionally, patched neurons were held for at least 15 minutes to permit biocytin filling for post-hoc morphological analysis and identification. Data were acquired via a HEKA EPC10USB amplifier and digitized at 20kHz in Patchmaster Next (HEKA). We did not correct for the liquid junction potential.

### Analysis of patch clamp recordings

Analysis of electrophysiological responses to hyperpolarizing and depolarizing current injections were performed using MATLAB and Python. In MATLAB, a custom script was used to import (Keine, 2022), standardize, and save the data as a .mat variable. A custom python script was used to import and analyze the data stored in the mat variable, partly using the electrophysiology feature extraction library (eFEL) (Van Geit et al., 2016) of the Blue Brain Project (Blue Brain Project, 2015). Briefly, eFEL was used to mark specific sweep events including the values and indices of action potential (AP) thresholds, peaks, and after hyperpolarizations (AHP) to calculate physiological parameters including half-height width, interspike interval, instantaneous firing frequency, and max firing frequency. The half-height width (HHW) of each AP (from AP threshold to AHP) was calculated by setting half the maximum amplitude ((Peak value - AP threshold value) / 2) as a horizontal threshold, interpolating a line containing 1000 points over the AP, identifying when the interpolated line crossed the threshold during the rising and falling phases, and subtracting the values from each other. The interspike interval (ISI) was calculated by subtracting the latter AP threshold value from the former (e.g., ISI[i] = APthres[i+1] - APthres[i]). The instantaneous firing frequency (IFF) was calculated by dividing the interspike interval values by 1 (e.g., IFF = 1/ISI). The max firing frequency was calculated by selecting the maximum frequency value (e.g., MFF = max (IFF)).

### Immunohistochemistry

After patch clamp recordings, ex vivo slices were stored in 4% PFA overnight. Free-floating sections were washed in 0.1M PBS and incubated with 1% NaBH4 for 20 minutes, washed again, and then incubated with 10% MeOH + 3% H2O2 for 15 minutes. Slices were then incubated in blocking buffer (5% NGS + 2% BSA + 0.5% triton X-100) for 1 hour at RT followed by overnight incubations of 1° antibodies (Chk α-GFP (1:1000) Rockland 600-901-215; Rb α-FLAG (1:1000) Sigma F7425) diluted in blocking buffer at RT. The following day, slices were washed with 0.1M PBS and incubated with 2° antibodies (Streptavidin-Cy5 (1:300) Jackson Immunoresearch 016-170-084; Goat α-Chk 488 (1:500) ThermoFisher Scientific A-11039; Goat α-Rb 594 (1:500) ThermoFisher Scientific A-11037) for 4 hours at RT. Slices were washed with PBS, mounted onto slides (ThermoFisher Scientific 12-550-15) via Aquamount (ThermoFisher Scientific, 13800) and cover-slipped (ThermoFisher Scientific 12-540-B) before imaging. For one FSI slice, we tested a fast-optical clearing method (FOCM) because it has been shown to enhance the imaging of fine structures, such as synaptic contacts, in thick sections with minimal loss of endogenous fluorescence and tissue distortion in mice (Zhu et al, 2019). We did not observe any significant distortion to the tissue or morphology of our biocytin filled neuron using this method and this cell was included in the analysis.

### Confocal imaging

Confocal Z-stack images were obtained using a Zeiss LSM 800 confocal microscope. Fixed slices containing filled neurons from ex vivo recordings were imaged using a 40x oil immersion objective (Plan-Apochromat 40x/1.4 Oil DIC (UV) VIS-IR M27) set at 1.0 airy disk unit. The biocytin filled cell was centered in the field of view, and the zoom factor was set to a minimum of 0.5x or adjusted to capture the entirety of the dendritic field. Images acquired at either 1024 x 1024 or 2048 x 2048 pixels with a 0.5x zoom factor corresponded to a minimum voxel dimension of 0.312 x 0.312 x 0.530 µm or 0.156 x 0.156 x 0.530 µm respectively in the X, Y, and Z dimensions. Fluorescence acquisition settings were as follows: Alexa Fluor 488 (excitation 493λ, emission 517λ, detection 490–550λ), Alexa Fluor 568 (excitation 577λ, emission 603λ, detection 565– 642λ), Alexa Fluor 647 (excitation 653λ, emission 668λ, detection 645–700λ). The pinhole size of the objective was set to 1.0 airy disk unit when capturing the longest wavelength fluorophore (Alexa Fluor 647). When imaging the other fluorophores (Alexa Fluor 405, 488, and 568), the pinhole size was adjusted to match the airy disk size of the 647 laser. This ensured that the scan with each laser excited fluorophores in the same sized area. Optical laser intensities were set individually for each channel and optimized throughout the Z-stack using the Z-stack auto-brightness correction tool. Z-stack ranges were set manually by tracking the range of the entire labeled cell.

### Morphological measurements

For each neuron in our dataset, we obtained the position along the AP, ML, and DV axes by manually aligning the coronal field view of the recorded slice with the corresponding Allen brain atlas figure (Allen Reference Atlas). We were not able to obtain the ML and AP position for one SPN and one FSI. Dendritic field measurements were obtained by creating a 2D orthogonal projection of the confocal Z-stack and then measuring the largest diameter of an elliptical sphere encompassing the neuron using ImageJ (Fiji) software. The range along the Z-axis for each cell was calculated as the distance between the first and last appearance of neuronal processes in the Z-stack.

### Imaris 3D-reconstructions of single neurons

Carl Zeiss image files were imported into Imaris version 9.7 equipped with the Filament Tracer plugin (Bitplane). Biocytin filled neurons were first reconstructed into 3D-surface objects using the “surfaces” tool. For our surface reconstructions, the “background subtraction” option was enabled and the “smoothing” option was disabled to avoid adding artificial curvature to the cell (Fogarty et al, 2013). We determined the largest diameter setting by measuring the largest cross-sectional diameter of the soma in the “slice view” mode. Using an interactive histogram of volumetric pixels, we manually adjusted the threshold to include as much of the neuron as possible while aiming to exclude any extraneous background fluorescence. Neurons were then reconstructed using the “filament tracer” plugin using a semi-automated tracing method. Reconstructing the neuron as a filament allowed us to segment the neuron into separate branch levels and dendrites, as well as obtain distance measurements for M1 and S1 inputs from the beginning of the filament (soma). To construct the neuron as a “filament” we manually traced the dendrites using the “autopath” tool. Once the final outline of the neuron was created, the diameter of the dendrites was adjusted using an automatic threshold. For SPNs, we used the automatic spine detection tool in the “filament” tab to detect and reconstruct dendritic spines. The diameter and length of spines were measured in “slice viewer” using min/max diameters of 0.6 µm and 2.5 µm for automatic spine detection thresholds. This step was excluded for our FSI dataset due to the lack of spines on their dendrites. We verified that there was a single filament for each reconstruction and that all the branch points and dendrites were accurately represented in comparison to the raw fluorescence images.

### M1 and S1 corticostriatal synaptic puncta identification and quantification

The Imaris “surface” reconstruction enabled us to mask the raw fluorescence from M1 and S1 channels onto the neuron. Using the “edit” tab within the surface reconstructed neuron, we masked putative presynaptic contacts from M1 and S1 corticostriatal projections by setting the intensity of voxels outside of the surface to 0. This allowed us to filter putative presynaptic contacts so that only fluorescence from M1 and S1 that colocalized with the surface reconstruction remained. This step resulted in two new fluorescent channels representing masked fluorescence from M1 and S1 onto the neuron.

Fluorescence from masked M1 and S1 inputs were made into 3-D spots using the “spots” tool. First, the cross-sectional diameter of putative presynaptic puncta from cortical projections was measured in “slice viewer”. The “different spot sizes” option was checked because fluorescent puncta from M1 and S1 varied in size. We chose a minimum diameter of 1µm and the “background subtraction” option was disabled. Spots were manually thresholded using an interactive histogram and then validated by verifying that the new spot object was representing masked fluorescence from a corticostriatal projection and not from extraneous background fluorescence. The “spots region type” was set to “local contrast” and the “spots regions” diameter threshold was set to “region border”. We used automatic thresholds for these two parameters and found that they gave us the best representation of our puncta as 3D-objects.

In Imaris, the default “spots close to filament” function is capable of filtering and quantifying M1 and S1 spots whose center lies within a specified distance from the edge of a neuron reconstructed as a filament object. However, it was limited in that it provided us with the absolute distance of each spot from the soma instead of considering the length of the dendrites. To circumvent this issue, we used a publicly available custom python script provided by Dr. Matthew Gastinger (Gastinger, 2022). Using the “spots close to filament” function provided by this script, we filtered S1 and M1 spots so that only those with edges were within 0.5 µm from the edge of the filament-constructed neuron remained, and we identified these spots as putative M1 and S1 inputs. This custom script also provided us with the quantity of inputs and the distance from the soma measured along the dendrite for each spot object.

### Statistical analyses

All statistical analyses were performed using GraphPad Prism software. For each metric, we first tested if the data was normally distributed using the Shapiro-Wilk test (Ghasemi & Zahediasl, 2012; Mishra et al, 2019). For normally distributed data, parametric comparative t-tests were used. A statistical F-test was first used to compare variances between groups. If the standard deviation between two normally distributed data sets was significantly different, the unequal variance Welch’s t-test was used. For unpaired samples that were not normally distributed the nonparametric Mann-Whitney U test was used. For paired data that was not normally distributed we used the Wilcoxon matched pairs signed rank test. For multiple group comparisons, we used the analysis of variance (ANOVA) statistical test. All data are presented as the mean +/- SEM.

### Code accessibility

A custom Imaris extension written and provided by Dr. Matthew Gastinger is publicly available through Github (Gastinger, 2022). Scripts used for the analysis of the electrophysiological recordings are available through the Margolis Github. (https://github.com/margolislab)

## Results

### M1 and S1 corticostriatal projections converge in the dorsolateral striatum

We injected adeno associated viruses (AAVs) encoding spaghetti monster fluorescent proteins (sm.FPs) GFP or Ruby into whisker M1 and whisker S1 to observe putative synaptic contacts onto striatal neurons via confocal light microscopy. We used sm.FPs because of their utility in multicolor experiments, enhanced fluorescence, and better resolution when imaging fine structures such as synaptic terminals (Viswanathan et al, 2015). To ensure there was no bias in the expression of sm.FPs due to the injection site, we counterbalanced the injection of sm.FPs in M1 and S1 throughout the experiments. In total, 7 mice were injected with sm.FP-GFP in S1 and sm.FP-Ruby in M1, and 3 mice were injected with sm.FP-GFP in M1 and sm.FP-Ruby in S1.

After waiting at least 3 weeks to permit viral expression, coronal slices containing the anterior striatum were obtained. Ex-vivo whole-cell patch clamp recordings were performed ipsilateral to the injection site to characterize the electrophysiological properties and fill individual neurons with biocytin (Fig 1A). Recorded and filled neurons were located within extensively overlapping sm.FP-labeled M1 and S1 corticostriatal projections in the dorsolateral region of the striatum (DLS) (Fig. 1B). Our final data set includes 14 striatal neurons from 10 mice (2 males, 8 females). 13 additional cells from 6 mice were excluded from reconstruction and analysis due to incomplete biocytin cell fills, inconclusive immunohistochemistry containing artifacts, excessive background fluorescence, or the inability to be identified as an SPN or FSI via their morphology or electrophysiology.

**Figure 1.**
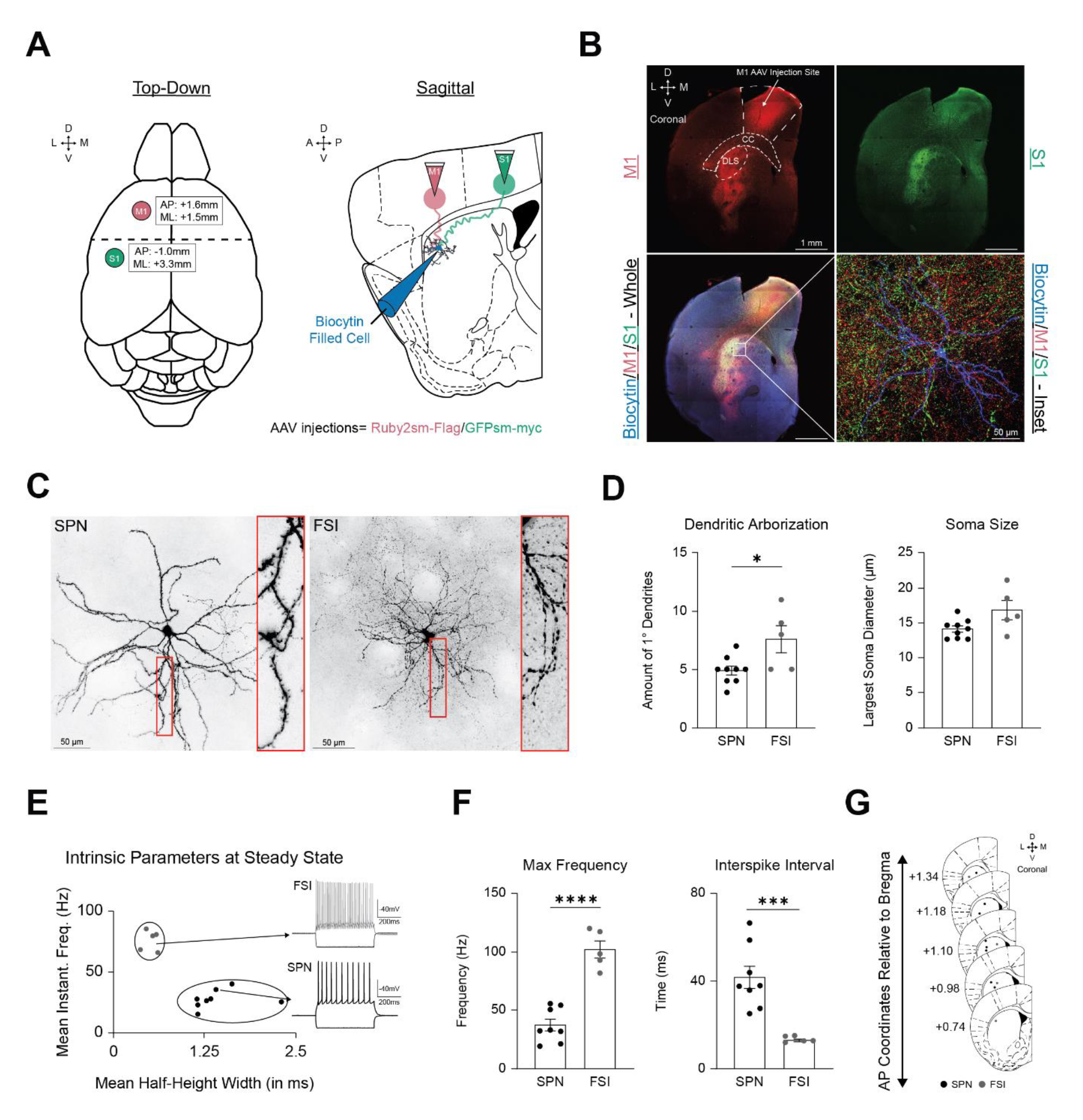
Viral circuit mapping of M1 and S1 corticostriatal projections to dorsal striatal SPNs and FSIs. **(A)** Viral circuit mapping strategy highlighting spaghetti monster injections into M1 and S1 from a top-down view (left) and the corticostriatal projection pattern onto biocytin filled striatal cells from a sagittal view (right). **(B)** *Top-left*: Low magnification view of a coronal striatal section containing the M1 sm.FP injection site and corticostriatal projections to the dorsolateral striatum. *CC =* corpus callosum, *DLS* = dorsolateral striatum. *Top-Right*: Low magnification view of a coronal striatal section with S1 corticostriatal projections labeled with sm.FPs. *Bottom-Left*: Low magnification view of merged fluorescence from S1 and M1 corticostriatal projections innervating a striatal SPN filled with biocytin (white box). *Bottom-Right*: High magnification inset bottom-left image with corticostriatal innervation by M1 and S1 onto a biocytin filled SPN. **(C)** *Left:* Pseudo colored representative image of a biocytin filled SPN with spiny dendrites (inset). *Right:* Pseudo colored representative image of a biocytin filled FSI with aspiny dendrites (inset). **(D)** *Left:* Average number of primary dendrites (note that FSIs have more primary dendrites extending from the soma compared to SPNs). *Right:* Average cross-sectional diameter of the soma **(E)** Comparison of mean instantaneous firing frequency and mean half-height width at steady state permits differentiation of FSIs from SPNs due to their fast firing rates. **(F)** *Left:* Average max firing frequency. *Right:* Average interspike interval. **(G)** Schematic representation of the recording location of the SPNs and FSIs in our dataset (note that all cells were recorded in the anterior dorsal striatum where M1 and S1 innervation was dense).

**Figure 1-1.**
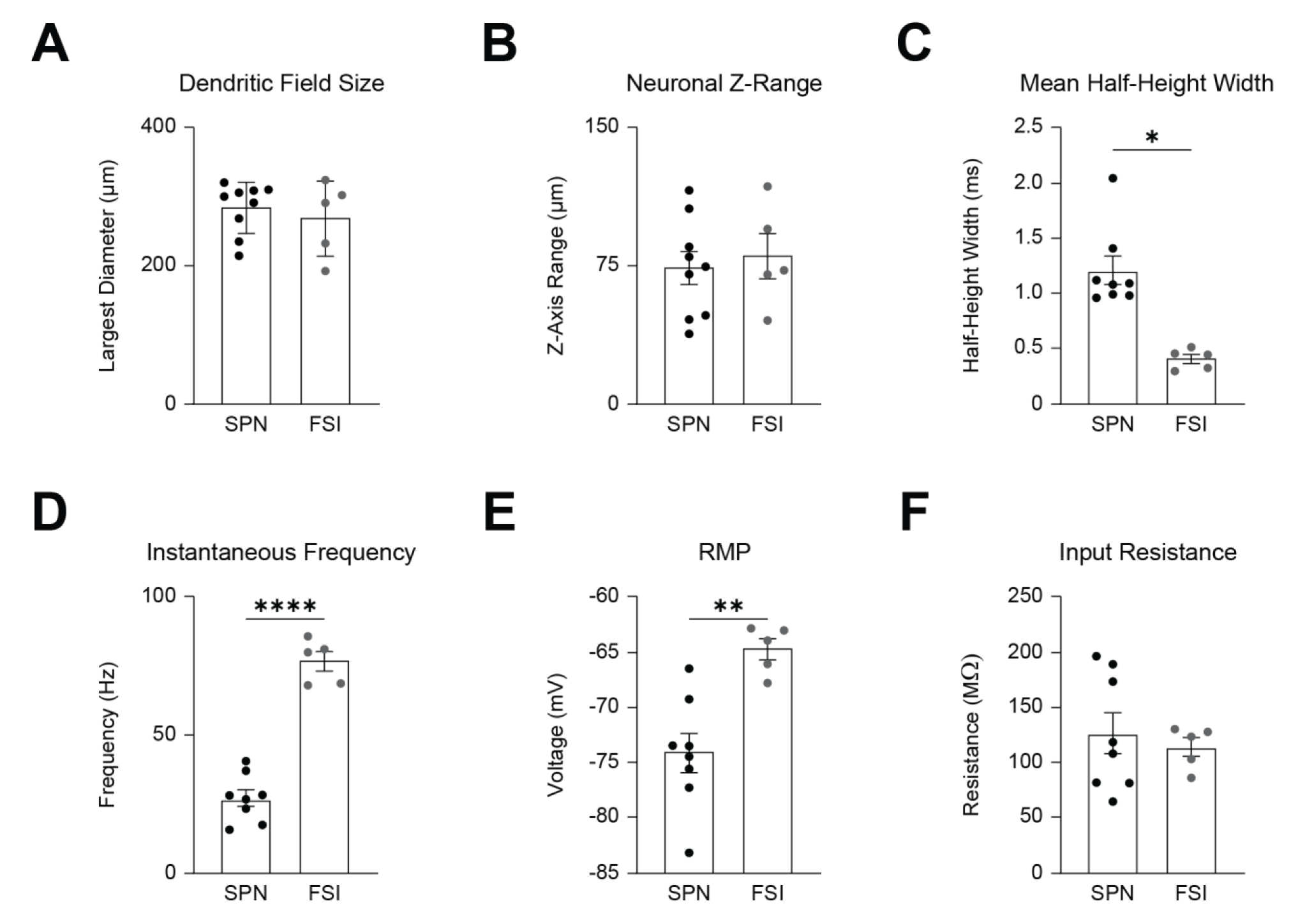
Morphological and electrophysiological properties of SPNs and FSIs. **(a)** Mean cross sectional diameter of the dendritic field. **(b)** Mean depth of the dendritic field along the z-axis. **(c)** Mean half height width (HWH) amplitude. **(d)** Mean instantaneous firing frequency. **(e)** Mean resting membrane potential voltage. **(f)** Mean input resistance.

### Patched SPNs and FSIs are identified by their distinct morphological and electrophysiological properties

Striatal neurons were patched without the use of genetic cell identification due to the expression of sm.FPs GFP and Ruby in cortical inputs, which limited our ability to record from SPNs or FSIs labeled with fluorescent markers such as GFP or TdTomato. Instead, patched striatal neurons were identified by their morphological and electrophysiological features, as in previous studies (Kawaguchi, 1993 & 1997; Tepper et al, 2018). We identified two distinct populations of striatal neurons in our dataset. Since 95% of the striatal population consist of SPNs, there was a high probability of patching them relative to FSIs (Hjorth et al, 2020). These experiments did not distinguish between D1- and D2-SPNs.

Most of the population (N= 9 cells) was identified as SPNs primarily based on the presence of prominent spines embedded along the dendrites (Fig. 1C). In comparison, a second population (N= 5 cells), identified as FSIs, often contained a fusiform-shaped soma, frequently branched, varicose aspiny dendrites, and dense axonal arborizations surrounding the dendritic field (Fig. 1C). We found no significant differences in the maximum size of the dendritic field (SPN = 284.6 +/- 12.39µm, N = 9 vs FSI = 269.1 +/- 24.52µm, N = 5, p = 0.539, t = 0.6323, df = 12, unpaired two tailed t-test) (Fig. 1-1a) or the range of the dendrites along the Z-axis between the two groups of neurons (SPN = 73.81 +/- 8.974µm, N = 9 vs FSI = 80.34 +/- 12.38µm, N = 5 p = 0.6748, t = 0.4301, df = 12, unpaired two tailed t-test) (Fig. 1-1b). However, SPNs had significantly fewer primary dendritic branches (SPN = 4.88 +/- 0.3889, N = 9 vs FSI = 7.6 +/- 1.166, N = 5, p = 0.0183, t = 2.728, df = 12, unpaired two tailed t-test) and slightly smaller soma diameter compared to FSIs (SPN = 14.26 +/- 0.4528µm, N = 9 vs FSI = 16.98 +/- 1.389µm, N = 5, p = 0.1228, t = 1.865, df = 7.206, unequal variance Welch’s t-test) (Fig. 1D), which is similar to what has been previously reported (Kawaguchi, 1993 & 1997; Fino & Venance, 2011; Tepper et al, 2018).

FSIs are characterized by their higher frequency (up to 100 Hz) action potential firing and shorter action potential duration, thus we could compare the mean instantaneous frequency and the mean half-height width of action potentials evoked in current clamp recordings to distinguish them from other types of aspiny neurons in the striatum (Kawaguchi, 1993 & 1997; Tepper et al, 2018) (Fig. 1E). The subpopulation of aspiny neurons identified as FSIs had shorter mean half-height widths (FSI _HHW_ = 0.40 +/- 0.04 ms, N = 5 vs SPN _HHW_ = 1.21 +/- 0.13 ms, n = 8, p = 0.0016, Mann-Whitney U = 0) (Fig. 1E, 1-1c), faster mean instantaneous firing frequencies (FSI_IFF_ = 76.84 +/- 3.56 Hz, N = 5 vs SPN _IFF_ = 26.99 +/- 3.07 Hz, n = 8, p < 0.0001, t = 10.38, df = 11, unpaired two tailed t-test) (Fig. 1E, 1-1d), more depolarized resting membrane potentials (FSI _RMP_ = -64.69 +/- 0.960 mV, N = 5 vs SPN _RMP_ = -74.16 +/- 1.787 mV, n = 8 vs, p = 0.0024, t = 3.924, df = 11, unpaired two tailed t-test) (Fig. 1-1e), faster maximal firing frequencies (FSI _MFF_ = 102.3 +/- 7.348 Hz, N = 5 vs SPN _MFF_ = 37.02 +/- 5.146 Hz, n = 8, p < 0.0001, t = 7.505, df = 11, unpaired two tailed t-test) (Fig. 1F), and shorter interspike intervals (FSI _ISI_ = 13.27 +/- 0.6246 ms, N = 5 vs SPN = 41.68 +/- 5.137 Hz, n = 8, p = 0.0008, t = 5.490, df = 7.206, unequal variance Welch’s t-test) (Fig. 1F) compared to SPNs.

SPNs and FSIs had similar input resistances (FSI _Input Resistance_ = 114.0 +/- 8.513 MΩ, N = 5 vs SPN _Input Resistance_ = 126.6 +/- 18.65 MΩ, n = 8, p = 0.6217, t = 0.5077, df = 11, unpaired two tailed t-test) (Fig. 1-1f). One neuron did not reach steady state firing during patch clamp recordings and was excluded from the analysis of ephys parameters but was still included in the SPN group based on its dendritic morphology.

To verify that there were no differences in the recording site within the striatum for SPNs and FSIs, we determined the position of the recorded cell relative to the anatomical features of the slice for a subset of neurons. Most neurons were obtained in the dorsal aspect of the striatum where M1 and S1 corticostriatal projections were concentrated and there were no significant differences in the recording location for both types of neurons (SPN _ML_ = + 1.738 +/- 0.059, n = 8 vs FSI _ML_ = +1.700 +/- 0.2245mm, n = 4, p = 0.8808, t = 0.1614, df = 3.430, unequal variance Welch’s t-test; SPN _AP_ = + 1.120 +/- 0.0414, n = 8 vs FSI _AP_ = + 1.040 +/- 0.1249, n = 4, p = 0.4555, t = 0.7763, df = 10, unpaired two tailed t-test; SPN _DV_ = -2.265 +/- 0.1101, n = 8 vs FSI _DV_ = -2.125 +/- 0.1199, n= 4, p = 0.4516, t = 0.7833, df = 10, unpaired two tailed t-test) (Fig. 1G) (Table 1).

**Table 1.** Position of recorded striatal neurons from ex-vivo slices relative to bregma. The medial lateral (ML), anterior posterior (AP), and dorsal ventral (DV) position of the recorded neuron within the striatum relative to bregma.

|  | ML | AP | DV |
| --- | --- | --- | --- |
| SPN, *n* = 8 | 1.738 +/- 0.06mm | 1.12 +/- 0.04mm | -2.265 +/- 0.10mm |
| FSI, *n* = 4 | 1.70 +/- 0.19mm | 1.04 +/- 0.11mm | -2.125 +/- 0.10mm. |

Overall, the neurons in our dataset have morphological and electrophysiological properties that are consistent with the well-documented differences between SPNs and FSIs. Using the absence of spines on the dendrites and the presence of fast spiking enabled us to differentiate between FSIs and SPNs without the use of genetically encoded fluorescent markers, such as parvalbumin, which is commonly used to label FSIs (Kawaguchi, 1993; Tepper et al, 2018).

### Corticostriatal inputs to SPNs and FSIs reconstructed in 3D using Imaris

The strength of synaptic connectivity from M1 and S1 to striatal SPNs and FSIs may derive from differences in the quantity and distribution of synaptic inputs. Based on our previous work (Lee et al. 2019), we hypothesized that SPNs have significantly more synaptic inputs from M1 compared to S1, but FSIs have a similar number of synaptic inputs from M1 and S1. To test for this, we quantified M1 and S1 inputs onto SPNs and FSIs from confocal 3D reconstructions using Imaris, similar to previous methods (Fogarty et al, 2013; Kuljis et al, 2019).

In brain slices from mice with M1 and S1 already labeled with sm.FPs, we patched and filled striatal neurons with biocytin for anatomical analysis. Raw fluorescence from single cell filled SPNs and FSIs was first converted into 3D “surface” objects using the surface tool in Imaris. Then, we constructed our neurons as “filament” objects because this enabled us to segment the dendritic spines, branch points, terminals, and reduce the inclusion of background fluorescence in the reconstruction (Fig. 2A).

**Figure 2.**
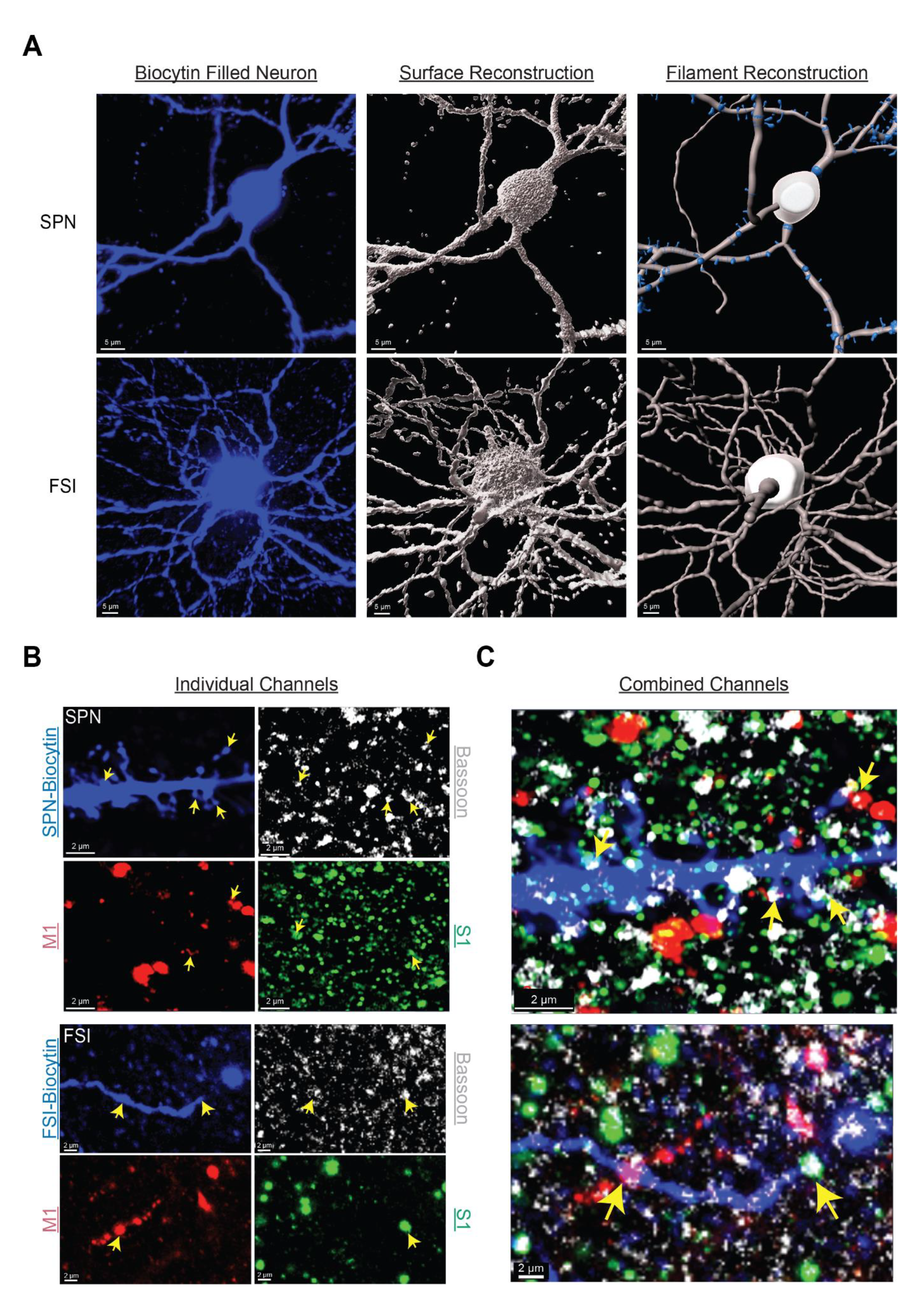
3D reconstructions of striatal neurons and confirmation of S1 and M1 synaptic puncta. **(A)** 3D-reconstruction of biocytin filled SPNs and FSIs into a “surface” and “filament” object in Imaris. **(B)** Representative raw fluorescence from biocytin, the presynaptic protein bassoon, M1, and S. **(C)** Merged image from **B** demonstrating S1 and M1 presynaptic inputs dually innervating an SPN (top) and FSI (bottom).

The dual labeling of M1 and S1 permitted visualization of both cortical inputs simultaneously, but the distance between the pre and postsynaptic membrane is beyond the limitations of confocal light microscopy (Maidorn et al, 2016). To verify that sm.FPs from M1 and S1 represent putative synaptic inputs, we captured fluorescence from M1 and S1 axons along with a marker for the presynaptic scaffolding protein, bassoon, in a subset of patched neurons. Puncta from sm.FPs that labeled M1 and S1, colocalized with bassoon along the dendrites and spines of filled neurons for both SPNs and FSIs (Fig. 2B,C). However, the small size of presynaptic puncta (<0.5µm), combined with the large depth of our Z-stack (>50µm), limited our ability to capture inputs colocalized with bassoon in deeper parts of the Z-stack because of significant photobleaching of bassoon.

Therefore, to reconstruct M1 and S1 inputs onto striatal neurons, raw fluorescence from corticostriatal puncta was first masked onto the surface of the reconstructed neuron and then converted into 3D “spot” objects. We quantified the number and distribution of spots whose edge was within ≤0.5µm from the edge of the filament construction and identified them as putative presynaptic inputs (Fig 3A). To ensure that our counts were not limited to one side of the Z-stack or confounded by limited light penetration, the Z positions of the first and last appearance of a dendrite, the soma, as well as the Z positions of identified M1 and S1 inputs were obtained throughout the stack. We confirmed that there were no inputs counted outside the range of the neuron within the Z-stack and that M1 and S1 fluorescence penetrated equally throughout the stack for both cell types (Fig. 3-1a).

**Figure 3.**
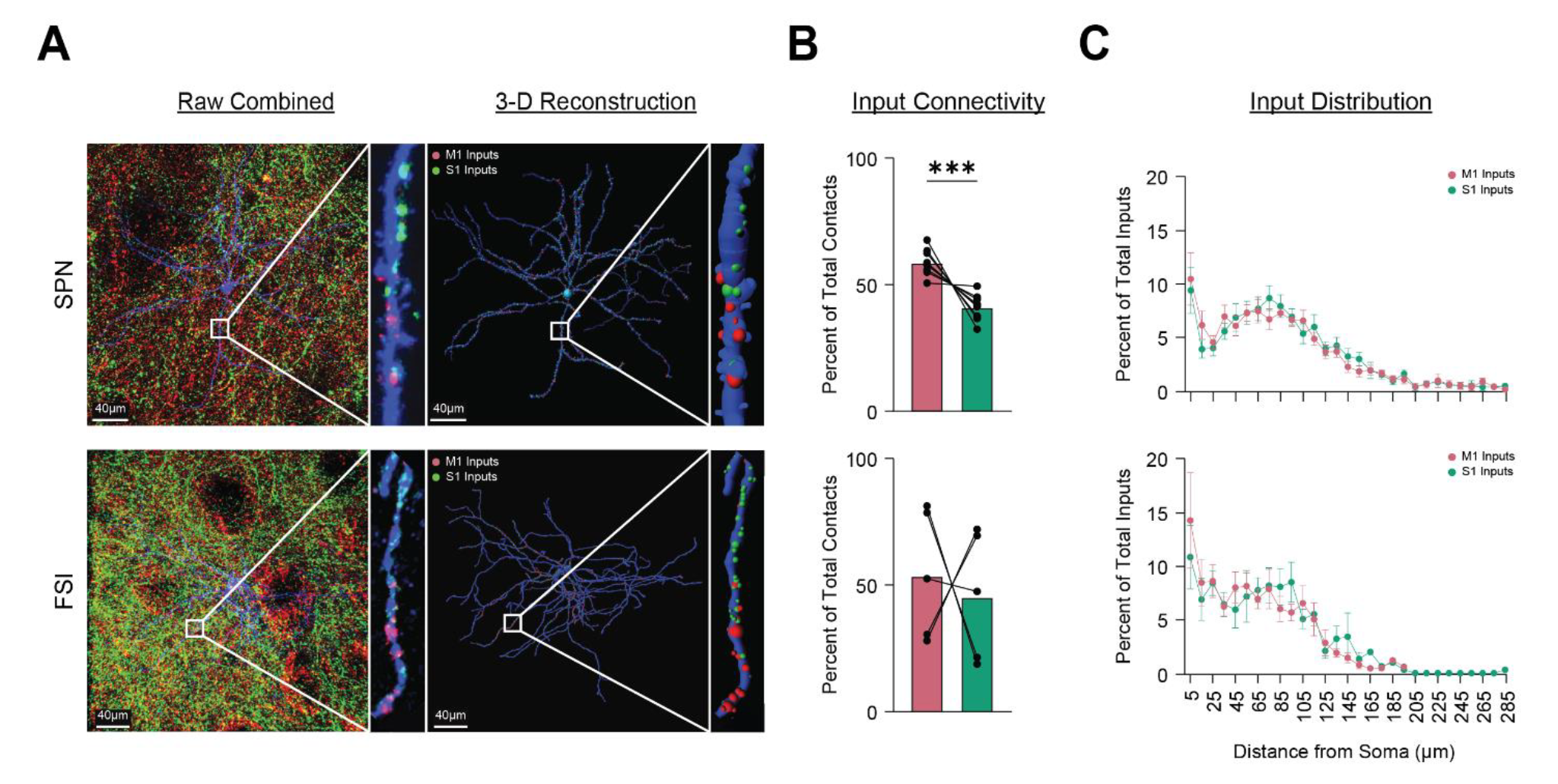
Differential innervation by S1 and M1 to SPNs and FSIs. (**A)** Magnified view of raw fluorescence (left) from S1 and M1 corticostriatal projections onto a biocytin filled SPN (top) and FSI (bottom) with their associated 3D-reconstructions (right). **(B)** Average percent innervation by M1 and S1. **(C**) Mean distribution of S1 and M1 inputs across the neuron when measuring the distance from the soma along the length of a dendrite.

**Figure 3-1.**
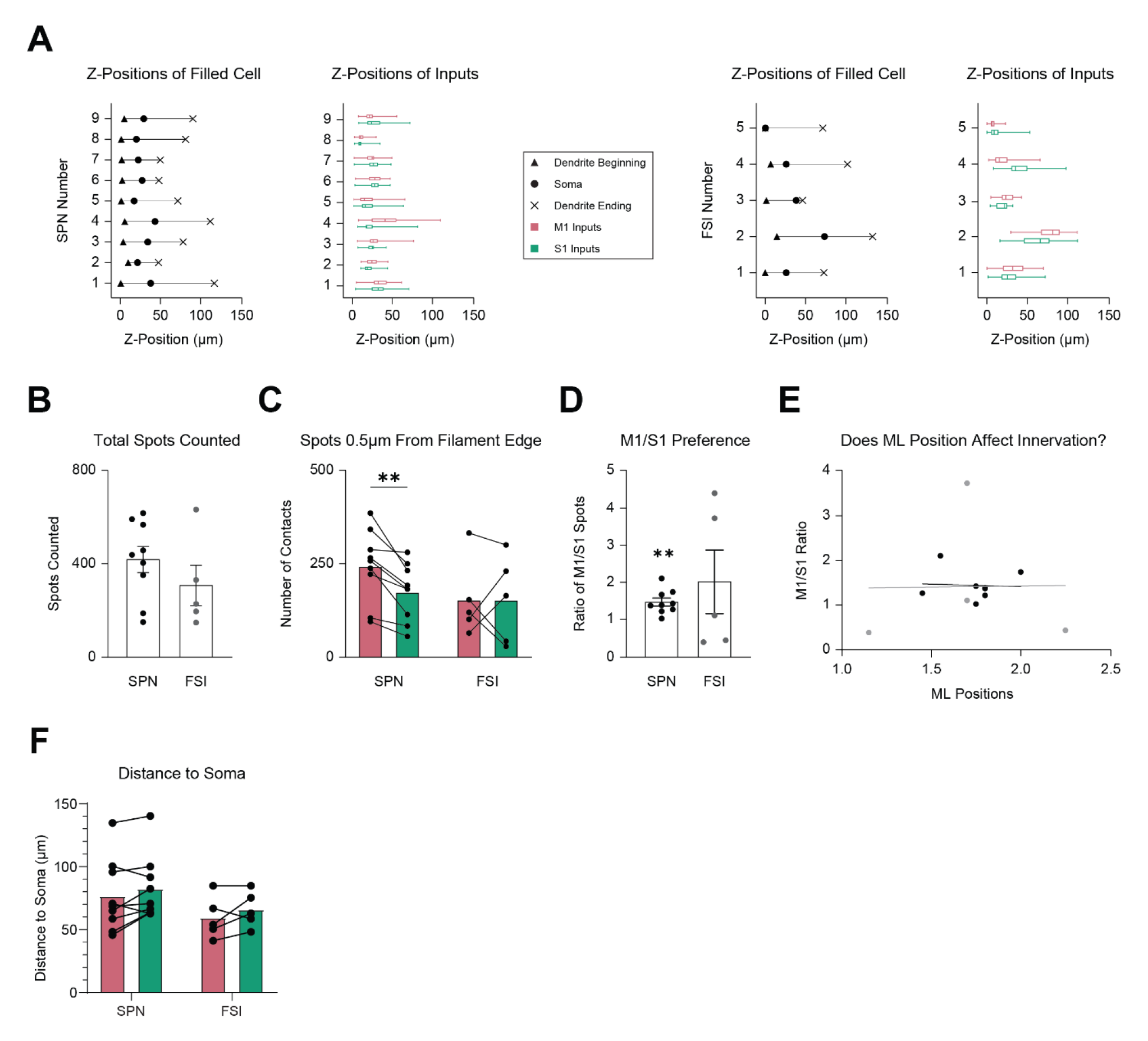
Distribution properties of S1 and M1 inputs to SPNs and FSIs. **(A)** Z-positions of the first and last appearance of a dendrite, and the soma of reconstructed SPNs (Left) and FSIs (Right) compared with the Z positions of their associated S1 and M1 inputs **(B)** Mean total number of combined S1 and M1 spots counted within 0.5µm of the filament constructed SPN or FSI **(C)** Mean number of spots counted from S1 and M1 within 0.5µm of the filament edge **(D)** Mean ratio of M1 spots to S1 spots counted within 0.5µm of the filament edge **(E)** Comparison of M1/S1 ratio with the mediolateral position of the neuron it was recorded from; black dots = SPNs (n=8), gray dots = FSIs (n=4). **(F)** Mean distance of an S1 and M1 input from the soma when measured along the length of the dendrite.

### Preferential anatomical innervation of SPNs by M1 compared to S1 corticostriatal inputs

On average, the total number of combined M1 and S1 inputs onto SPNs was greater than FSIs but not significantly different (SPN = 418.55 +/- 52.97, N = 9 vs FSI = 306.2 +/- 78.11, N = 5, P = 0.2797, t = 1.132, df = 12, unpaired two tailed t-test), suggesting that we counted a similar number of inputs within a given Z-stack for both cell types (Fig. 3-1b). However, there were significantly more M1 than S1 inputs onto SPNs (SPN_M1_ = 244.44 +/- 29.10 vs SPN_S1_ = 174.111 +/- 32.53, N = 9, p = 0.0025, t = 4.339, df = 8, paired two tailed t-test) (Fig. 3-1c), resulting in a considerably larger proportion of M1 than S1 inputs to SPNs (SPN_M1_ = 58.89 +/- 1.61% vs SPN_S1_ = 41.11 +/- 1.61%, N = 9, p = 0.0008, t = 5.222, df = 8, paired two tailed t-test) (Fig. 3B). The ratio of M1 inputs to S1 inputs on SPNs was significantly greater than a value of 1, indicating a stronger preference for the M1 input (SPN_M1/S1_ = 1.467 +/- 0.102, N = 9, p = 0.0025, t = 4.338, df = 8, one sample t-test) (Fig. 3-1d).

In comparison, there were no substantial difference between the number and proportion of M1 and S1 inputs onto FSIs (FSI_M1_ = 153.6 +/- 42.17 vs FSI_S1_ = 152.6 +/- 35.39, N = 5, p = 0.9849, t = 0.02018, df = 4, paired two tailed t-test) (FSI_M1_ = 54.19 +/- 10.25% vs FSI_S1_ = 45.81 +/- 10.25%, N = 5, p = 0.7332, t= 0.3656, df = 4, paired two tailed t-test) (Fig. 3B, 3-1c). Similarly, the ratio of M1 to S1 inputs was not significantly different from a value of 1, suggesting no overall bias for M1 or S1 (FSI_M1/S1_ = 2.013 +/- 0.766, N = 5, p = 0.302, t = 1.183, df = 4, one sample t-test) (Fig. 3-1d).

Neither the cell type (F (1,12) = 1.282, p = 0.2797, two-way repeated measures ANOVA) nor the origin of the cortical input (F (1,12) = 2.884, p = 0.1152, two-way repeated measures ANOVA) had a significant effect on the variability of the number of inputs counted from M1 or S1. In addition, there was no significant interaction between the effects of cortical input and cell type on the number of inputs counted (F (1,12) = 2.725, p= 0.1247, two-way repeated measures ANOVA). However, a significant source of variability on counts stemmed from individual subject variability (F (12,12) = 5.582, p = 0.0028, two-way repeated measures ANOVA) (Fig 3C). M1 and S1 projection fields can vary along the mediolateral axis of the striatum, however we found no significant correlation between the ratio of M1 to S1 inputs and the mediolateral position of recorded neurons within the DLS (Fig 3-1e), leaving the source of individual variability unknown.

Altogether, although M1 and S1 inputs dually innervate SPNs and FSIs, M1 provides a greater number of synaptic inputs than S1 to SPNs, whereas M1 and S1 provide overall equal input to FSIs. These results complement previous measures of the functional synaptic strength of these inputs (Lee et al, 2019).

### M1 and S1 inputs are similarly distributed in the proximal, medial, and distal regions of SPNs and FSIs

The spatial organization of synaptic inputs along the dendritic tree, in addition to the total number of inputs, is an important factor that can influence synaptic integration. To test if there are differences in the spatial distribution of M1 and S1 inputs to SPNs and FSIs, inputs were segmented based on their distance along the dendrites relative to the soma into 10µm bins. In both SPNs and FSIs, the distribution of M1 and S1 inputs closely followed each other and were found in proximal (0 < 30µm), medial (30 < 100µm), and distal regions (>100µm) of the neuron. Interestingly, the largest concentration of inputs was found near the soma for both SPNs and FSIs (SPN_M1 0-10µm_ = 10.50 +/- 2.453%, SPN_S1 0-10µm_ = 9.433 +/- 2.181% N = 9; FSI_M1 0-10µm_ = 14.31 +/- 4.473%, FSI_S1 0-10µm_ = 10.88 +/- 2.981%, N = 5) (Fig. 3C). This was followed by a sharp dip in the number of inputs synapsing 20-30µm away from the soma of the SPNs, but this was not as prominent in FSIs. The number of M1 and S1 inputs peaked again at distances 70-80µm from the soma in SPNs but not in FSIs, and then decreased to near zero values at distal dendritic locations >100µm from the soma in both cell types (Fig. 3C).

Overall, M1 and S1 inputs were distributed similarly with no significant differences in the mean distance of M1 and S1 inputs from the soma in both SPNs and FSIs (SPN_M1-Soma_ = 76.38 +/- 9.586µm vs SPN_S1-Soma_ = 82.36 +/- 8.527µm, N = 9, p = 0.2031, Wilcoxon matched pairs signed rank test; FSI_M1-Soma_ = 59.44 +/- 7.555µm vs FSI_S1-Soma_ = 65.97 +/- 6.427µm, N = 5, p = 0.2682, paired two tailed t-test) (Fig. 3-1f).

### M1 and S1 inputs cluster onto SPNs but not FSIs

Since individual striatal neurons receive both M1 and S1 inputs, we wanted to know whether these inputs are localized near each other. To determine if M1 and S1 inputs form synaptic clusters, we measured the distance between inputs of the same type (M1-M1 or S1-S1) or inputs of different types (M1-S1). In SPNs, M1 inputs were found in proximity (∼5µm) to other M1 inputs, and S1 inputs were also near other S1 inputs (SPN_M1-M1_ = 4.390 +/- 0.5693µm vs SPN_S1-S1_ = 4.532 +/- 0.4742µm, N = 9, p = 0.6728, t = 0.4383, df = 8, paired two tailed t-test). In FSIs, the shortest distance between M1-M1 inputs or S1-S1 inputs was similar to each other, but slightly larger than SPNs (FSI_M1-M1_ = 6.680 +/- 1.220µm vs FSI_S1-S1_ = 7.794 +/- 1.692µm, N = 5, p = 0.5832, t = 0.5961, df = 4 paired two tailed t-test) (Fig. 4A-C).

**Figure 4.**
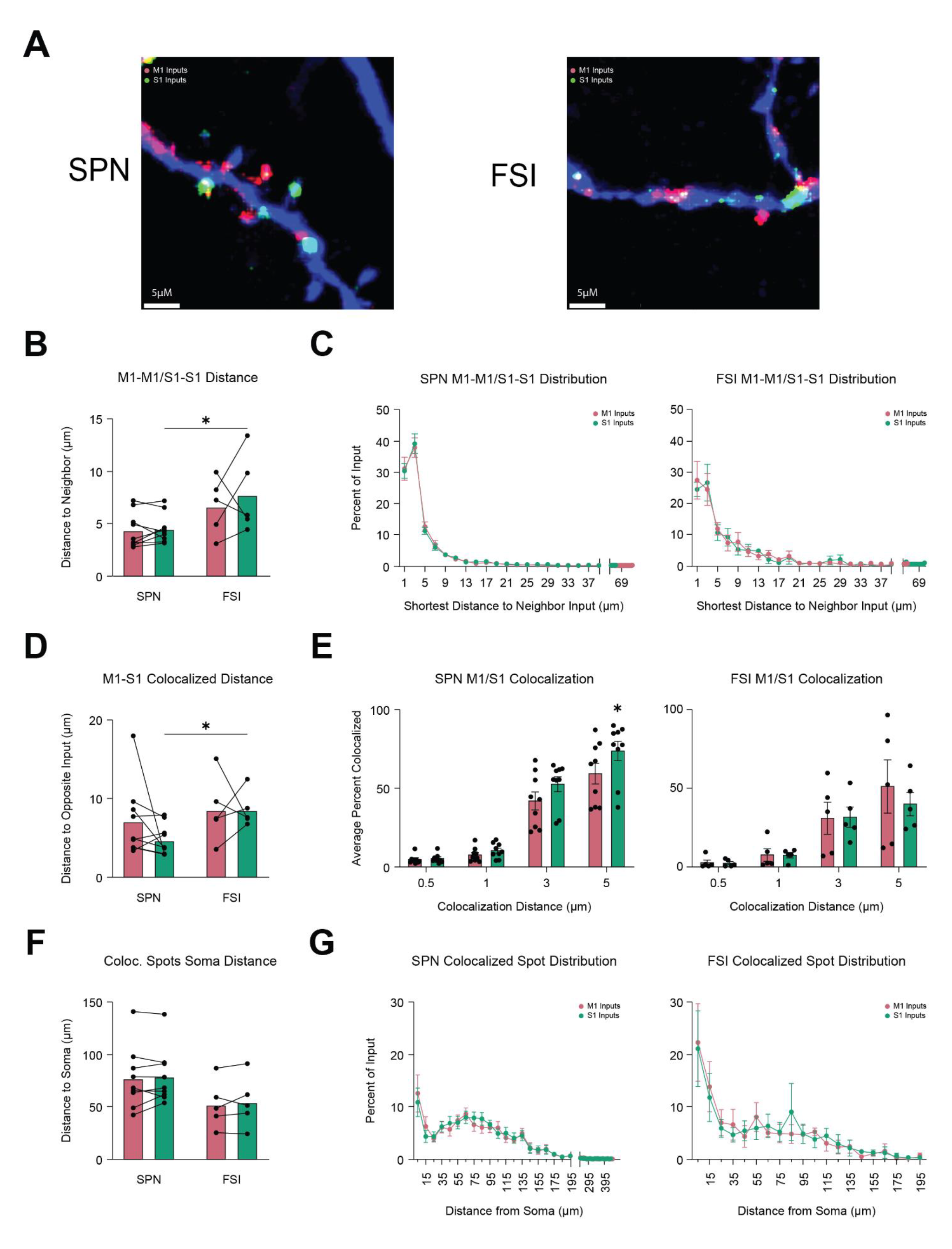
S1 and M1 input cluster on SPNs but not in FSIs. **(A)** Representative image of fluorescence from M1 (red) and S1 (green) masked onto the dendrites of an SPN (Left) and an FSI (Right). **(B)** Mean distance between the nearest input from the same cortical region. **(C)** Mean distribution of the shortest distance to the nearest input from the same cortical region. **(D)** Mean distance between inputs from different cortical regions. **(E)** Mean percent of S1 and M1 Inputs that colocalized with an increasing distance threshold for SPNs (Right) and FSIs (Left). **(F)** Mean distance from the soma when measured along the length of the dendrite for S1 and M1 spots that colocalized within 5µm of each other. **(G)** Distribution of colocalized spots from **F**.

The cell type had a statistically significant effect on the variability of the distance between inputs from the same cortical region (F (1,12) = 6.768, p = 0.0232, two-way repeated measures ANOVA). Therefore, we compared between cell types and found that S1 inputs in SPNs were significantly closer together than in FSIs (SPN_S1-S1_ = 4.532 +/- 0.4742 µm, N = 9 vs FSI_S1-S1_ = 7.794 +/- 1.692µm, N = 5, p = 0.0358, t = 2.364, df = 12 unpaired two tailed t-test) (Fig. 4B). A similar trend was observed for M1 inputs but was not significant (SPN_M1-M1_ = 4.390 +/- 0.5693µm, N = 9 vs FSI_M1-M1_ = 6.680 +/- 1.220µm, N = 5, p =0.0747, t = 1.592, df = 12 unpaired two tailed t-test) (Fig 4B). These results indicate that inputs from the same cortical region tend to cluster more on SPN dendrites compared to FSI dendrites.

We next asked whether inputs from different cortical regions were more clustered in SPNs than FSIs. We found that the mean distance between an M1 and S1 input was significantly smaller in SPNs than in FSIs (SPN_S1-M1_ = 4.727 +/- 0.6643µm, N = 9 vs FSI_S1-M1_ = 8.609 +/- 1.023µm, N = 5, p = 0.0061, t = 3.321, df = 12, unpaired two-tailed t-test; SPN_M1-S1_ = 7.156 +/- 1.565µm, N = 9 vs FSI_M1-S1_ = 8.605 +/- 1.906µm, N = 5, p = 0.6309, Mann-Whitney U = 18.50) (Fig. 4D). There was a greater chance of S1 inputs being found within 5 µm of an M1 input in SPNs (SPN_S1-5µm_ = 73.92 +/- 5.86%, p = 0.0049, t = 3.847, df = 8, SPN_M1-5µm_ = 59.44 +/- 6.23%, p = 0.1912, t = 1.428, df = 8, N = 9, one sample t-test) (Fig. 4E). This was not the case for FSIs, as less than 50% of the S1 input colocalized within 5 µm of an M1 input (FSI_M1-5µm_ = 51.18 +/- 15.33 %, p = 0.9483, t = 0.0689, df = 4, FSI_S1-5µm_ = 39.86 +/- 6.71%, p = 0.2478, t = 1.352, df = 4, N = 5, one sample t-test) (Fig. 4E). These results indicate that the majority of S1 inputs onto SPNs form synaptic clusters with M1, and that clustering is more prominent in SPNs compared to FSIs.

These results led us to examine the spatial distribution of M1 and S1 inputs that form clusters within 5µm of each other. We observed that clustered M1-S1 inputs were distributed across the proximal, medial, and distal regions for both cell types and there was no differences in the mean distance from the soma for inputs that were part of a cluster (SPN_M1-S1_ = 77.04 +/- 9.943µm, SPN_S1-M1_ = 78.76 +/- 8.807µm, N = 9; FSI_M1-S1_ = 52.11 +/- 10.37µm, FSI_S1-M1_ = 54.37 +/- 11.15µm, N = 5) (Fig. 4F). Similar to the distribution of individual M1 and S1 inputs, the highest concentration of clustered inputs was found 0-10µm from the soma (SPN _M1 0-10µm_ = 12.71 +/- 3.572, SPN_S1 0-10µm_ = 10.99 +/- 2.747, N = 9; FSI_M1 0-10µm_ = 22.51 +/- 7.452%, FSI_S1 0-10µm_ = 21.32 +/- 7.298%, N = 5) (Fig. 4G), with a second peak 60-70µm from the soma. The second peak was less prominent in FSIs compared to SPNs. Altogether, our results indicate that clustered M1 and S1 inputs are distributed across all regions of SPNs and FSIs but are located less densely at distal dendrites.

## Discussion

In this study, using dual fluorescent tracing, we measured the extent of corticostriatal projections from M1 and S1 to individual neurons in the DLS and identified cell-specific differences in the quantity and distribution of M1 and S1 inputs onto SPNs and FSIs. Our findings indicate that SPNs receive significantly more inputs from M1 compared to S1, while FSIs show no overall bias in the number of M1 or S1 inputs (Fig. 3). In addition, we found that M1 and S1 inputs have similar distributions across proximal, medial, and distal regions of SPNs and FSIs, but in SPNs, they are found near each other, forming synaptic clusters (Fig. 4). These results have implications for how M1 and S1 corticostriatal inputs modulate striatal circuitry and behavior.

### Cell-specific differences in the number of inputs from M1 and S1 suggest stronger S1 connectivity to FSIs in the DLS

The convergence of M1 and S1 inputs to the DLS reflects its important role in sensorimotor integration (Makino et al, 2016; Gritton et al, 2019; Lipton et al, 2019; Matiros et al 2019). We observed extensive overlap of M1 and S1 projections onto individual neurons in the DLS, in agreement with previous experiments (Fig. 1B, Fig. 2B, C) (Hoffer & Alloway, 2001; Hunnicutt et al, 2016; Hooks et al, 2018).

Due to our use of biocytin to label patched neurons, we did not use genetic labels for D1 and D2 SPNs, or parvalbumin for FSIs. However, in our study the proportion of inputs from M1 and S1 to SPNs had low variability (Fig 3C), suggesting that D1 and D2 SPNs have a similar distribution of M1 and S1 inputs, which we previously demonstrated through optogenetic stimulation of M1 and S1 inputs (Lee et al, 2019). The more notable differences in M1 and S1 innervation were found between SPNs and FSIs, and we distinguished patched FSIs from SPNs by their larger soma size, varicose aspiny dendrites, and high frequency spiking compared to SPNs, which has been reported in previous studies (Fig 1C-F, Fig. 1-1) (Kawaguchi, 1993 & 1997; Tepper et al, 2018). While most parvalbumin interneurons in the striatum are FSIs, there are likely regional subtypes of FSIs with distinct responses and connectivity to cortical brain regions (Monteiro et al, 2018; Tokarska & Silberberg, 2022). Future experiments could use further genetic cell typing, which could help clarify the variability we observed between individual cells.

SPNs and FSIs require many excitatory inputs to generate action potentials due to their hyperpolarized resting membrane voltages seen in Fig. 1-1e (Kawaguchi, 1993 & 1997; Tepper et al, 2018). A greater quantity of inputs leads to increased EPSPs because inputs can spatially summate on dendrites (Magee, 2000). M1 and S1 can form synaptic contacts onto the same SPN or FSI (Fig. 2B) (Ramanathan et al, 2002; Johansson & Silberberg, 2020; Charpier et al, 2020), but DLS SPNs have weak responses to sensory stimuli and strong responses to motor activity (Lee et al, 2019; Matiros et al, 2018, Charpier et al, 2020). In contrast, FSIs respond to M1 and S1 stimulation with EPSPs of similar amplitude (Lee et al, 2019; Johansson & Silberberg, 2020). Directly comparing the number of inputs from M1 or S1 revealed no significant differences between SPNs and FSIs, however we found that SPNs had significantly fewer inputs from S1 compared to M1, and that number of inputs between M1 and S1 was similar in FSIs (Fig 3-1c). The reduced innervation by S1 compared to M1 in SPNs, but equal innervation compared to M1 in FSIs suggest that S1 promotes the feedforward inhibition of SPNs by preferentially activating FSIs over SPNs (Lee et al 2019, Johansson and Silberberg 2020).

A large source of variability in the counts stemmed from our group of FSIs (Fig 3-1c, see Anova results). However, this is not due to differences in the penetration of light from our sm.FP constructs during imaging (Fig. 3-1a), or the total number of inputs counted between SPNs and FSIs (Fig. 3-1b). Moreover, despite a denser overlap of M1 and S1 in the more lateral aspects of the striatum we found no major correlation between lateral position and the ratio of M1/S1 inputs. This might indicate that ratio of M1 to S1 inputs to SPN and FSIs is consistent across the DLS (Fig. 1G, Fig. 3-1e) instead of the possibility that the mediolateral position of a patched neuron biased innervation patterns towards M1 or S1.

### The location of M1 and S1 inputs onto SPNs and FSIs suggest integration through spatiotemporal mechanisms

The anatomical measures indicate that the quantity of synapses partly account for differences between cell types. The distance between the input from the soma is also important because distal inputs undergo electrotonic decay and result in smaller EPSPs recorded at the soma when compared to proximal inputs (Rall, 1967; Magee, 2000; Straub et al, 2016). We found that M1 and S1 inputs had similar distributions across regions of the neuron for both SPNs and FSIs. A large proportion of apparent inputs were observed closer to the soma for SPNs (Fig. 3C). However, reports using correlative light and electron microscopy (CLEM) with VGLUT1, asynaptic marker expressed in corticostriatal terminals, have also observed cortical terminals near the cell body but demonstrated that true asymmetric synapses form at the dendritic spines of SPNs, with few inputs to the dendritic shafts and cell body (Reiner et al, 2010; Lei et al, 2013, Deng et al, 2015).

In SPNs, spines are prominent on the primary dendrites distal to the first dendritic ramifications (Fig. 1C, 2A) (Wilson & Groves,1980), likely explaining why we observed a drop in counts in more proximal regions (Fig. 3C). Interestingly, the number of M1 and S1 inputs peaked again at medial dendritic distances (70-80µm) (Fig. 3C), where SPNs have active calcium conductances that contribute to the generation of “up-states” and increased excitability (Plotkin et al, 2011). Other reports have shown stronger synaptic responses in SPNs and the presence of synaptic clusters at similar dendritic locations (Straub et al, 2016; Hwang et al, 2022). In contrast, “up-states” have not been demonstrated in FSIs, suggesting that they integrate inputs passively (Plenz and Kitai, 1998); Our data is consistent with reports that corticostriatal synapses are broadly distributed onto the dendrites of FSIs (Ramanathan et al, 2002; Nakano et al, 2018; Zheng et al, 2021).

SPNs can perform sublinear, linear, and supralinear integration of excitatory inputs depending on the timing and location of the input (Carter et al, 2007, Straub et al, 2016). The clustering of synapses is optimized for long-term plasticity because the coincident activation of multiple synapses in spines that are in close together on a dendrite can produce a self-regenerating dendritic spike, which is stronger and longer lasting than an EPSP. (Losonczy & Magee, 2006; Carter et al, 2007; Kastellakis & Poirazi, 2019, Du et al, 2017). Although we observed less S1 inputs compared to M1 in SPNs (Fig. 3A, B), the majority (> 50%) of S1 inputs were in proximity (< 5µm) to a neighboring S1 input or M1 input, and this was not the case for FSIs (Fig. 4A-D). These results support the theory that SPNs can act as coincidence detectors for M1 and S1 and integrate signals from these cortical regions through temporal and spatial summation by clustering sensorimotor inputs. Critically, it has been observed that stimulation of M1 inputs induces an initial membrane depolarization in SPNs that is quickly enhanced by activity generated from S1 (Charpier et al, 2020), and M1 inputs from mice trained on motor tasks form active clusters on the spines of SPNs in the DLS (Hwang et al, 2022).

### Implications of findings on sensorimotor integration by SPNs and FSIs in the DLS

In vivo, during sensorimotor learning, SPNs form ensembles that are highly active during the initiation and termination of task-related movements associated with reward (Matiros et al, 2018; Gritton et al, 2019). PV-FSIs have an important contribution to SPN ensemble formation in the DLS (Matiros et al, 2018; Gritton et al, 2019). The cell specific distribution of M1 and S1 to the DLS that is seen in our results likely contributes to organizing which cells are active during this process.

Synaptic clustering greatly increases the likelihood that activity from M1 and S1 is coincidentally detected on short sections of the dendritic membrane, leading to large depolarizations and changes in plasticity (Carter al al, 2007; Kastellakis & Poirazi, 2019). This increases the chance that these inputs become associated with other cortical and thalamic regions when there is convergent activity (Carter et al, 2007; Huerta-Ocampo et al, 2014, Makino et al, 2016). Therefore, it is likely that while S1 alone does not evoke strong responses in SPNs, SPNs that receive convergent activity from task related M1 and S1 neurons will overcome local inhibition by FSIs and increase the probability of firing an action potential. As a result, they become part of task related circuits in the DLS that underlie habitual responses to sensory stimuli (Matiros et al, 2018; Lipton et al, 2019).

In conclusion, the organization of M1 and S1 inputs to the DLS suggest that activity from S1 will preferentially excite the striatal FSI population, leading to feed forward inhibition of SPNs during sensorimotor integration. Our findings have significant implications for how corticostriatal circuits encode learned movements. Future investigations should examine if there are cell-specific requirements for plasticity in corticostriatal inputs to SPNs and FSIs.

## Acknowledgements

This work was funded by the National Institutes of Health (NIH F31 NS124343) (NIH R01NS094450) and the National Science Foundation (NSF IOS-1845355)

